# Gene model for the ortholog of *tgo* in *Drosophila busckii*

**DOI:** 10.64898/2026.06.26.734908

**Authors:** Jorge Perez, Abbie A. Giunta, Jacqueline K. Wittke-Thompson

## Abstract

Gene model for the ortholog of *tango* (*tgo*) in the Sep. 2015 (UC Berkeley ASM127793v1/DbusGB1) Genome Assembly (GenBank Accession: GCA_001277935.1) of *Drosophila busckii*. This ortholog was characterized as part of a developing dataset to study the evolution of the Insulin/insulin-like growth factor signaling pathway (IIS) across the genus *Drosophila* using the Genomics Education Partnership gene annotation protocol for Course-based Undergraduate Research Experiences.

## Introduction

*This article reports a predicted gene model generated by undergraduate work using a structured gene model annotation protocol defined by the Genomics Education Partnership (GEP; thegep.org) for Course-based Undergraduate Research Experience (CURE). The following information in quotes may be repeated in other articles submitted by participants using the same GEP CURE protocol for annotating Drosophila species orthologs of Drosophila melanogaster genes in the insulin signaling pathway*.

In this GEP CURE protocol students use web-based tools to manually annotate genes in non-model *Drosophila* species based on orthology to genes in the well-annotated model organism fruit fly *Drosophila melanogaster*. The GEP uses web-based tools to allow undergraduates to participate in course-based research by generating manual annotations of genes in non-model species (Rele et al., 2023). Computational-based gene predictions in any organism are often improved by careful manual annotation and curation, allowing for more accurate analyses of gene and genome evolution (Mudge and Harrow 2016; Tello-Ruiz et al., 2019). These models of orthologous genes across species, such as the one presented here, then provide a reliable basis for further evolutionary genomic analyses when made available to the scientific community.” (Myers et al., 2024).

“The particular gene ortholog described here was characterized as part of a developing dataset to study the evolution of the Insulin/insulin-like growth factor signaling pathway (IIS) across the genus *Drosophila*. The Insulin/insulin-like growth factor signaling pathway (IIS) is a highly conserved signaling pathway in animals and is central to mediating organismal responses to nutrients (Hietakangas and Cohen 2009; Grewal 2009).” (Myers et al., 2024).

“The Drosophila *tango* (CG11987, FBgn0264075) gene encodes a bHLH-PAS protein that controls CNS midline and tracheal development (Sonnenfeld et al., 1997). Both cell culture and in vivo studies have shown that a DNA enhancer element acts as a binding site for both Single-minded::Tango and Trachealess::Tango heterodimers and functions in controlling CNS midline and tracheal transcription. Isolation and analysis of tango mutants reveal CNS midline and tracheal defects. In addition, the bHLH-PAS proteins Similar (Sima) and Tango (Tgo) function as HIF-α and HIF-β homologues, respectively, in a conserved hypoxia-inducible transcriptional response in *Drosophila melanogaster* that is homologous to the mammalian HIF-dependent response (Lavista-Llanos et al., 2002). Insulin has been shown to activate HIF-dependent transcription, both in *Drosophila* S2 cells and in living *Drosophila* embryos, and this effect is mediated by PI3K-AKT and TOR pathways. Overexpression of dAKT and dPDK1 in normoxic embryos provoked a major increase in Sima nuclear localization, mimicking the effect of a hypoxic treatment (Dekanty et al., 2005).” (Lawson et al., 2025).

“*D. busckii* (NCBI:txid30019) belongs to the *Dorsilopha* subgenus (Sturtevant 1942) of the genus *Drosophila* first described by Coquillett in 1901 (Accessed December 22, 2022 from the Integrated Taxonomic Information System

(ITIS), https://doi.org/10.5066/F7KH0KBK). The *Dorsilopha* subgenus is sibling to the *Sophophora* subgenus of *Drosophila* (Remsen and O’Grady 2002). *D. busckii*, possibly native to North America, is cosmopolitan with established populations around the world in primarily temperate regions (CABI Compendium, Accessed December 22, 2022), and has extremely broad host preferences (Sturtevant 1921).” (Backlund et al., 2025).

We propose a gene model for the ortholog in *D. busckii* of the *D. melanogaster* tango (*tgo)* gene. The genomic region of the ortholog corresponds to the uncharacterized protein XP_017847579.1 (Locus ID LOC108603353) in the Sep. 2015 (UC Berkeley ASM127793v1/DbusGB1) assembly of *D. busckii* (Zhou *et al*, 2015; GCA_001277935.1; PRJNA274996). This gene model is based on RNAseq data from *D. busckii* (Zhou *et al*, 2015; PRJNA274996) and *tgo* in *D. melanogaster using* FlyBase release FB2024_02 (GCA_000001215.4; Gramates et al., 2022; Jenkins et al., 2022; Larkin et al., 2021).

## Synteny

*tgo* occurs on chromosome 3R in *D. melanogaster* and is flanked by *hyx* and *neur* upstream and *CG11986* and *Sf3b5* downstream. We determined that the putative ortholog of *tgo* is found on scaffold CP012526.1 in *D. busckii* with LOC108603353 (via *tblastn* search with an E-value of 0.0 and percent identity of 73.88%), where it is surrounded by LOC108603354 (XP_017847583.1), LOC108603352 (XP_017847583.1), LOC108604002 (XP_017848731.1), and LOC108604004 (XP_017848732.1) which correspond to *hyx, neur, CG11986*, and *Sf3b5* in *D. melanogaster* with E-values and percent identities (0.0 and 91.68%; 0.0 and 77.02%; 0.0 and 74.10%; 7e-69 and 94.12%) respectively, as determined by *blastp* (Figure 1A, Altschul et al., 1990). We suggest this is the correct ortholog assignment for *tgo* in *D. busckii* because local synteny is conserved and the percent identity between tgo-PA in *D. busckii* and tgo-PA in *D. melanogaster* is high (81.0%).

**Figure 1.**
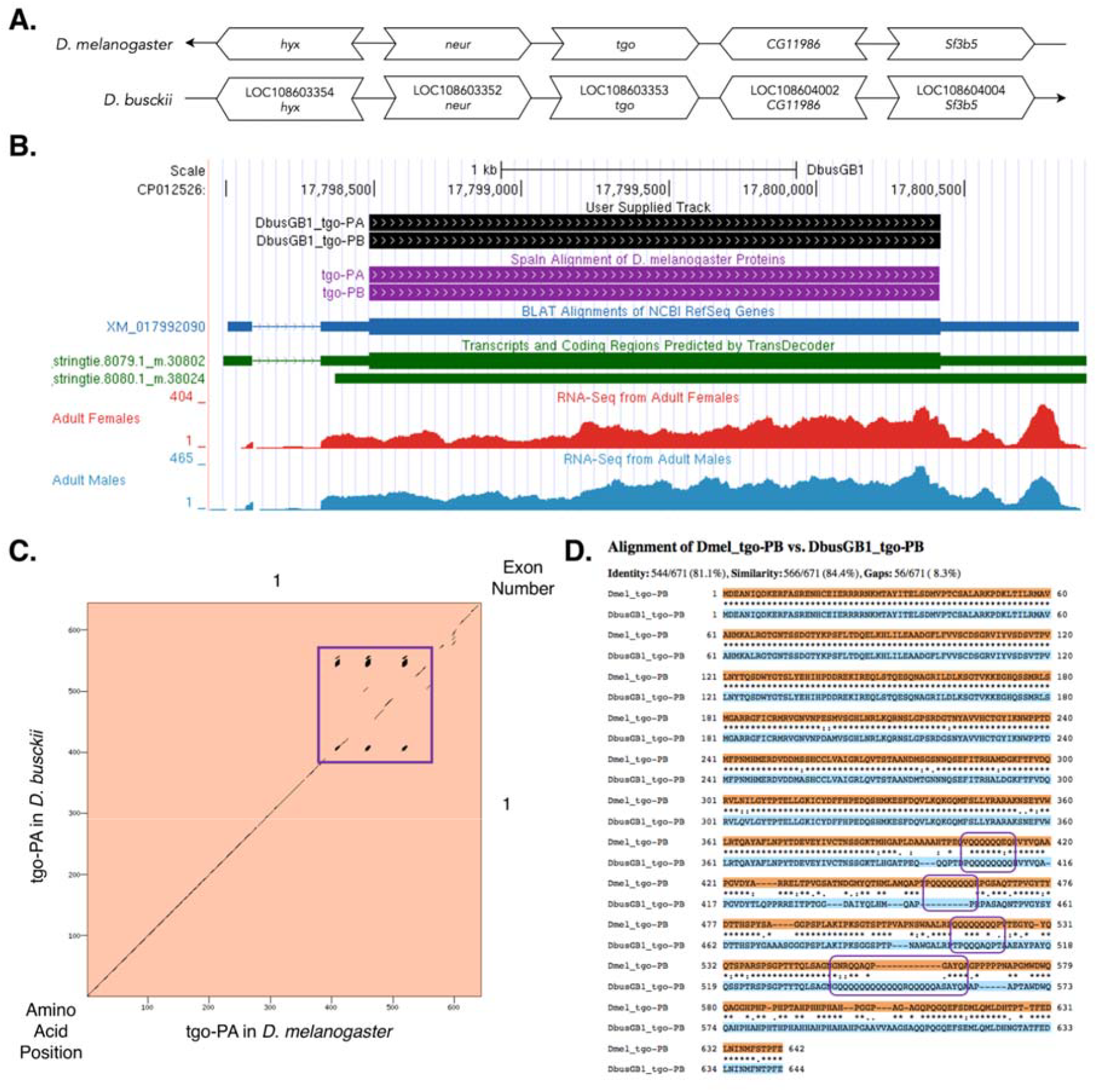
*tgo* gene model comparison between *Drosophila busckii* and *Drosophila melanogaster* orthologs. **(A)Synteny of genomic neighborhood of *tgo* in *D. melanogaster* and *D. busckii***. Gene arrows pointing in the same direction as *tgo* in both *D. busckii* and *D. melanogaster* are on the same strand as the target gene; gene arrows pointing in the opposite direction are on the opposite strand. The thin underlying arrow pointing to the right indicates that *tgo* is on the + strand in *D. busckii*; arrows pointing to the left indicate that *tgo* is on the – strand in *D. melanogaster*. White arrows in *D. busckii* indicate the locus ID and the orthology to the corresponding gene in *D. melanogaster*. The gene names given in the *D. busckii* gene arrows indicate the orthologous gene in *D. melanogaster*, while the locus identifiers are specific to *D. busckii*. **(B) Gene Model in UCSC Track Hub** (Raney et al. 2014): the gene model in *D. busckii* (black), Spaln of D. melanogaster Proteins (purple), alignment of refseq proteins from *D. melanogaster*), BLAT alignments of NCBI RefSeq Genes (blue, alignment of refseq genes for *D busckii*), RNA-Seq from Adult Females (red) and Adult Males (blue), alignment of Illumina RNAseq reads from *D. busckii*), and Transcripts (green) including coding regions predicted by TransDecoder and Splice Junctions Predicted by regtools using *D. busckii* RNA-Seq (Zhou *et al*, 2015; PRJNA274996). The custom gene model (User Supplied Track) is indicated in black with the exon depicted with a wide box (arrows indicate direction of transcription). **(C) Dot Plot of tgo-PA in *D. melanogaster* (*x*-axis) vs. the orthologous peptide in *D. busckii* (*y*-axis)**. Amino acid number is indicated along the left and bottom; exon number is indicated along the top and right, and exons are also highlighted with alternating colors. There are multiple instances of tandem repeats highlighted by the purple box. There are also a few indels represented by skewed lines. **(D) The protein alignment of tgo-PA in *D. melanogaster* against tgo-PA in *D. busckii***. The tandem repeats of Q (Glutamine) are contained within the purple boxes.

## Protein Model

*tgo* in *D. busckii* has two identical protein coding isoforms (tgo-PA and tgo-PB) which both contain one protein-coding exon (Figure 1B). Similarly, *tgo* in *D. melanogaster* has two identical protein coding isoforms (tgo-PA and tgo-PB) with one protein-coding exon. The sequence of tgo-PA in *D. busckii* has 76.42% identity with tgo-PA in *D. melanogaster* as determined by *blastp* (Figure 1C). There were multiple instances of repeats as demonstrated by dark black dots in the Dot Plot and contained within the purple boxes in the protein alignment (Figures 1C and 1D). This gene model can also be seen within the target genome at this <u>TrackHub</u>.

### Special characteristics of the protein model

**Tandem repeats and indel at the end of tgo-PA:** There are multiple instances of tandem repeats, specifically amino acid Q (Glutamine), near the end of the exon as highlighted by the purple boxes in the protein alignment. (Figure 1D) There are also a couple of indels demonstrated by the dashes in the protein alignment. A dash in the *D. melanogaster* line means a deletion in that species and an insertion in *D. busckii*. A dash in the *D. busckii* line indicates a deletion in that species and an insertion in *D. melanogaster* (Figure 1D).

## Methods

“Detailed methods including algorithms, database versions, and citations for the complete annotation process can be found in Rele et al. (2023). Briefly, students use the GEP instance of the UCSC Genome Browser v.435 (https://gander.wustl.edu; Kent et al., 2002; Navarro Gonzalez et al., 2021) to examine the genomic neighborhood of their reference IIS gene in the *D. melanogaster* genome assembly (Aug. 2014; BDGP Release 6 + ISO1 MT/dm6). Students then retrieve the protein sequence for the *D. melanogaster* reference gene for a given isoform and run it using *tblastn* against their target *Drosophila* species genome assembly on the NCBI BLAST server (https://blast.ncbi.nlm.nih.gov/Blast.cgi; Altschul et al., 1990) to identify potential orthologs. To validate the potential ortholog, students compare the local genomic neighborhood of their potential ortholog with the genomic neighborhood of their reference gene in *D. melanogaster*. This local synteny analysis includes at minimum the two upstream and downstream genes relative to their putative ortholog. They also explore other sets of genomic evidence using multiple alignment tracks in the Genome Browser, including BLAT alignments of RefSeq Genes, Spaln alignment of *D. melanogaster* proteins, multiple gene prediction tracks (e.g., GeMoMa, Geneid, Augustus), and modENCODE RNA-Seq from the target species. Detailed explanation of how these lines of genomic evidenced are leveraged by students in gene model development are described in Rele et al. (2023). Genomic structure information (e.g., CDSs, intron-exon number and boundaries, number of isoforms) for the *D. melanogaster* reference gene is retrieved through the Gene Record Finder (https://gander.wustl.edu/~wilson/dmelgenerecord/index.html; Rele et al., 2023). Approximate splice sites within the target gene are determined using *tblastn* using the CDSs from the *D. melanogaste*r reference gene. Coordinates of CDSs are then refined by examining aligned modENCODE RNA-Seq data, and by applying paradigms of molecular biology such as identifying canonical splice site sequences and ensuring the maintenance of an open reading frame across hypothesized splice sites. Students then confirm the biological validity of their target gene model using the Gene Model Checker (https://gander.wustl.edu/~wilson/dmelgenerecord/index.html; Rele et al., 2023), which compares the structure and translated sequence from their hypothesized target gene model against the *D. melanogaster* reference gene model. At least two independent models for a gene are generated by students under mentorship of their faculty course instructors. Those models are then reconciled by a third independent researcher mentored by the project leaders to produce the final model. Note: comparison of 5’ and 3’ UTR sequence information is not included in this GEP CURE protocol.” (Gruys et al., 2025)

## Supporting information

Gene model data files

## Supplemental Files

1.Zip file containing a FASTA, PEP, GFF files for the gene model

2.Figure 1 in high resolution

### Metadata

Bioinformatics, Genomics, *Drosophila*, Genotype Data, New Finding

## Acknowledgements

We would like to thank Wilson Leung for developing and maintaining the technological infrastructure that was used to create this gene model and Laura K. Reed for overseeing the project. We would also like to thank Aiden Long and Abigail Myers for their work reconciling this model. Thank you to FlyBase for providing the definitive database for *Drosophila melanogaster* gene models. Further, we would like to thank the editors and developers at the journal *microPublication: Biology* for assistance in developing the template for these single gene ortholog publications.

